# Bifunctional Architecture Enables Substrate Catalysis and Channeling in *Paracoccus* TMAO Demethylase

**DOI:** 10.1101/2025.11.26.690739

**Authors:** Trung Thach, KanagaVijayan Dhanabalan, Shiwangi Maurya, Yu Han-Hallet, Senwei Quan, Jane Allison, Gurunath Ramanathan, Ramaswamy Subramanian

## Abstract

Substrate channeling enhances efficiency and prevents toxicity by directing unstable intermediates between active sites. Trimethylamine N-oxide demethylase (TDM) degrades trimethylamine N-oxide (TMAO) to dimethylamine and formaldehyde (HCHO), but the fate of HCHO has remained unclear. We report cryo-EM structures of TDM in apo, substrate-, and product-bound states that reveal a previously unknown channeling pathway. Combined structural, biochemical, and target molecular dynamics analyses show that HCHO is generated in a catalytic core and guided through a tunnel to a remote tetrahydrofolate (THF)-binding site, where it forms methylene-THF. Thus, TDM emerges as a bifunctional enzyme that unites TMAO demethylation with one-carbon transfer, providing a mechanistic explanation for its role in metabolic efficiency and detoxification.

## INTRODUCTION

Substrate channeling occurs in enzymatic reactions, where intermediates are transferred directly from one enzyme’s active site to another, bypassing their release into the cellular environment.^[1,2]^ This mechanism improves catalytic efficiency by minimizing intermediate loss, while simultaneously protecting cells from the damaging effects of unstable or toxic species.^[1,2,3]^ Classic examples include glycolytic and amino acid metabolic enzymes, where covalent or noncovalent tunnels ensure safe and rapid passage of intermediates. ^[1,2,3]^ Despite its importance, structural evidence for substrate channeling remains limited, particularly in enzymes that couple detoxification with energy metabolism.

Trimethylamine (TMA) and its oxidized derivative trimethylamine N-oxide (TMAO) are abundant small molecules present in both marine organisms and the human gut microbiome.^[4,5]^ TMA arises from microbial metabolism of dietary choline, carnitine, and other animal-derived nutrients, and is subsequently oxidized to TMAO by host flavin-containing monooxygenases.^[4],[5]^ These metabolites are physiologically and clinically significant; elevated TMAO levels have been associated with cardiovascular disease and gastrointestinal cancer.^[4,5]^ Conversely, in marine bacteria and certain gut microbes, TMAO serves as both an energy source and a carbon donor, fueling specialized catabolic pathways that convert it into downstream metabolites.^[4,5,6]^

A key enzyme in this process is trimethylamine N-oxide demethylase (TDM), which catalyzes the cleavage of a methyl group from TMAO to generate dimethylamine (DMA) and formaldehyde (HCHO).^[6,7]^ Early biochemical studies revealed that TDM is a two-domain protein: an N-terminal domain that mediates TMAO demethylation, and a C-terminal domain structurally related to tetrahydrofolate (THF)-dependent enzymes, including the glycine cleavage T protein and dimethylsulfoniopropionate demethylase.^[7,8,9]^ This homology led to the hypothesis that the C-terminal domain could utilize the HCHO intermediate to form methylenetetrahydrofolate (MTHF), thereby coupling detoxification with one-carbon transfer metabolism.^[7,9]^ However, the molecular details of this bifunctional activity, how reactive formaldehyde is channeled and presented to the THF-binding site, have remained unresolved.

Here, we report cryo-EM structures of *Paracoccus* TDM captured in apo, substrate-bound, and product-bound states, providing the first high-resolution view of its dual-domain architecture (local resolution of 2.0 Å). Structural analysis revealed that the TMAO- and THF-binding pockets are separated by approximately 60 Å, raising the question of how formaldehyde is safely transferred. Tunnel mapping combined with target molecular dynamics simulations identified a continuous, negatively charged channel linking the two active sites, suggesting a structural basis for substrate channeling. Complementary biochemical assays and mass spectrometry confirmed that HCHO generated at the Core catalytic site is efficiently converted to MTHF at the C-terminal domain. Together, these results establish TDM as a bifunctional enzyme that unites TMAO demethylation with formaldehyde detoxification and one-carbon transfer, highlighting a microbial strategy to enhance metabolic efficiency while mitigating toxicity.

## RESULTS and DISCUSSION

### Optimization of TDM Activity and Evidence for Bifunctionality

We first optimized purification and assay conditions for TDM by systematically varying salt concentration, pH buffer, and temperature. These analyses revealed a broad range, with the highest activity between pH 6.0 and 7.0 at 150 mM NaCl and 35-45 °C (Figure S1A–D). Under these conditions, TDM robustly demethylated TMAO, yielding HCHO with a *V*_*max*_ of 156 nmol/min/mg and a *K*_*m*_ of 1.33 mM (Figure 1A). Notably, these kinetic parameters indicate higher catalytic activity than the previously reported TDM from *Methylocella silvestris*, which exhibits a *K*_*m*_of 3.88 mM and a *V*_*max*_of 14.61 nmol/min/mg.^[7]^ HPLC-MS confirmed the production of DMA as a co-product (Figure 1B), consistent with prior reports.^[7]^ Notably, HCHO concentrations decreased upon addition of THF (Figure 1C), suggesting that HCHO may be further processed into MTHF (Figure 1D). Overall, these findings suggest that TDM-mediated TMAO demethylation generates HCHO, which can subsequently react with THF, potentially linking TMAO breakdown to one-carbon metabolism.

**Figure 1.**
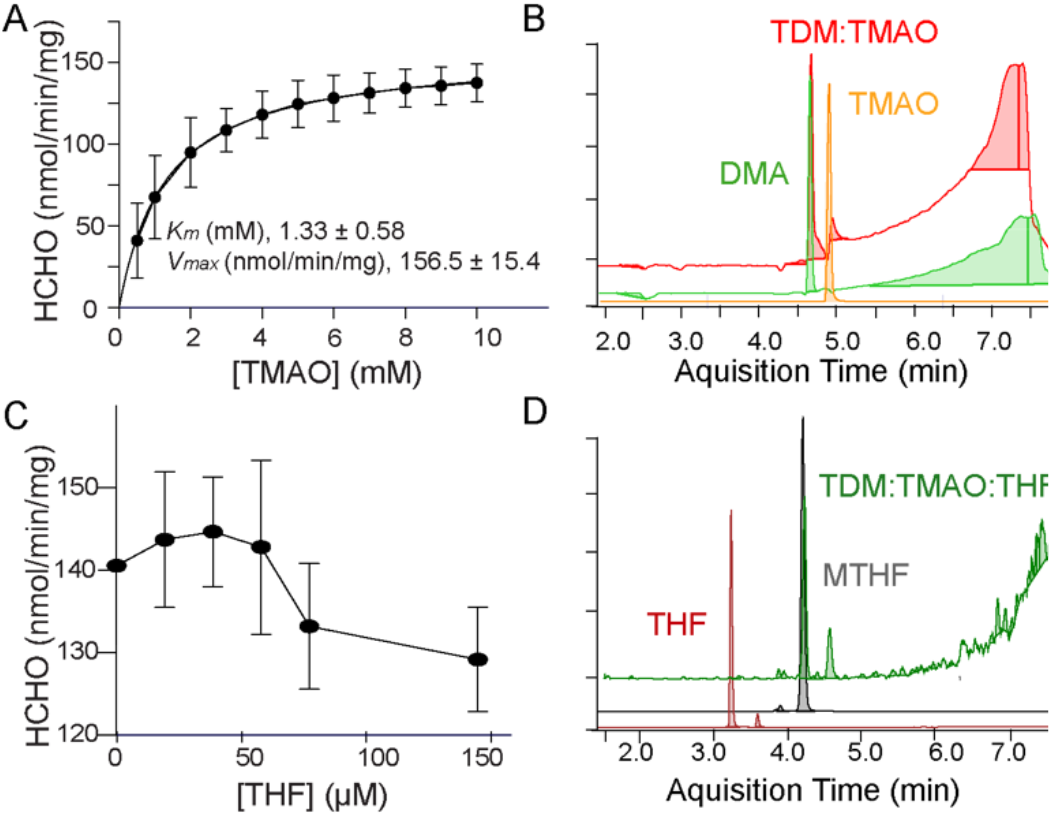
TDM catalyzes two sequential reactions. (A) Steady-state kinetic analysis of TDM - producing HCHO, in response to varying concentrations of the substrate TMAO (n=3). (B) Representative HPLC-MS spectrum of DMA product from TDM:TMAO reaction. The standard curve for DMA is shown in green, and that for TMAO is shown in orange. The red trace shows the product of the reaction when TMAO is added to TDM, indicating the formation of DMA. Counts in the Y-axis. (C) Linear regression analysis demonstrates a concentration-dependent decrease in HCHO production when THF is included in the reaction mixture, suggesting THF-dependent consumption or conversion of HCHO. (D) Representative HPLC-MS confirming the formation of MTHF adduct as a product of the TDM:TMAO:THF reaction. The red, gray, and green lines represent the standard curves for THF, MTHF, and the reaction product when TMAO and THF were present with TDM, respectively.

### Cryo-EM Structures Reveal an Unusual Oligomeric Assembly

To elucidate the catalytic mechanism, we determined cryo-EM structures of apo TDM, a product-bound state (TMAO-soaked wild type), and a substrate-trapped mutant (D220A/D367A in complex with TMAO) (Figure 2A–B, S2, Table 1). TDM assembles into an unexpected 2 + 2½ oligomeric architecture (Figure 2A–C). Two full-length subunits form a highly symmetric core dimer (RMSD = 0.12 Å over 787 Cα atoms), while two peripheral half-subunits consist solely of C-terminal THF-binding domains (Figure 2C). The core dimer interface is stabilized by extensive hydrogen-bonding and hydrophobic interactions (Figure S3).

**Table 1.** Cryo-EM data collection, refinement, validation and statistics.

| Structures | TDM-apo | TDM:DMA:HCHO:THF | (DD):TMAO:THF |
| --- | --- | --- | --- |
| PDB | 9Q59 | 9Q5L | 9Q5A |
| EMD | EMD-72223 | EMD-72239 | EMD-72224 |
| *Data collection and processing* |  |  |  |
| Magnification | 105,000 | 105,000 | 105,000 |
| Voltage (kV) | 300 | 300 | 300 |
| Electron exposure ((e<sup>-</sup>/Å<sup>2</sup>)) | 56.8 | 56.8 | 56.8 |
| Defocus range (μm) | 0.8-2.0 | 0.8-2.0 | 0.8-2.0 |
| Raw pixel size ( Å) | 0.411 | 0.411 | 0.411 |
| Symmetry imposed | C2 | C2 | C2 |
| Number of initial particle images | 145,414 | 568,305 | 240,010 |
| Number of final particle images | 26,866 | 174,871 | 143,707 |
| Map resolution (Å) | 2.76 | 2.79 | 2.80 |
| FSC threshold | 0.143 | 0.143 | 0.143 |
| Map resolution range ( Å) | 2.5-5.0 | 2.5-5.0 | 3.0-5.0 |
| *Refinement* |  |  |  |
| Initial model used | Alphafold model | Apo model | Apo model |
| Model resolution ( Å) | N/A | 2.8 | 2.8 |
| FSC threshold | 0.5 | 0.5 | 0.5 |
| Model resolution range ( Å) | N/A | 3.0-50 | 3.0-50 |
| Model composition |  |  |  |
| Non-hydrogen atoms | 17878 | 17058 | 17917 |
| Protein residues | 2309 | 2196 | 2306 |
| Ligand: ZN | 2 | 2 | 2 |
| TMO |  |  | 2 |
| HCHO |  | 2 |  |
| DMN |  | 2 |  |
| B factors (Å<sup>2</sup>) |  |  |  |
| Protein | 118.75 | 87.46 | 86.93 |
| Ligand | 144.67 | 89.10 | 90.75 |
| RMSD values |  |  |  |
| Bond lengths ( Å) | 0.004 | 0.004 | 0.003 |
| Bond angles (°) | 0.586 | 0.608 | 0.501 |
| *Validation* |  |  |  |
| Molprobity score | 2.22 | 2.32 | 2.02 |
| Clash score | 8.74 | 12.12 | 7.78 |
| Poor rotamer (%) | 2.98 | 2.63 | 2.18 |
| Ramachandran plot (%) |  |  |  |
| Favored | 94.25 | 93.89 | 95.07 |
| Allowed | 5.49 | 6.01 | 4.80 |
| Outliers | 0.26 | 0.09 | 0.13 |

**Figure 2.**
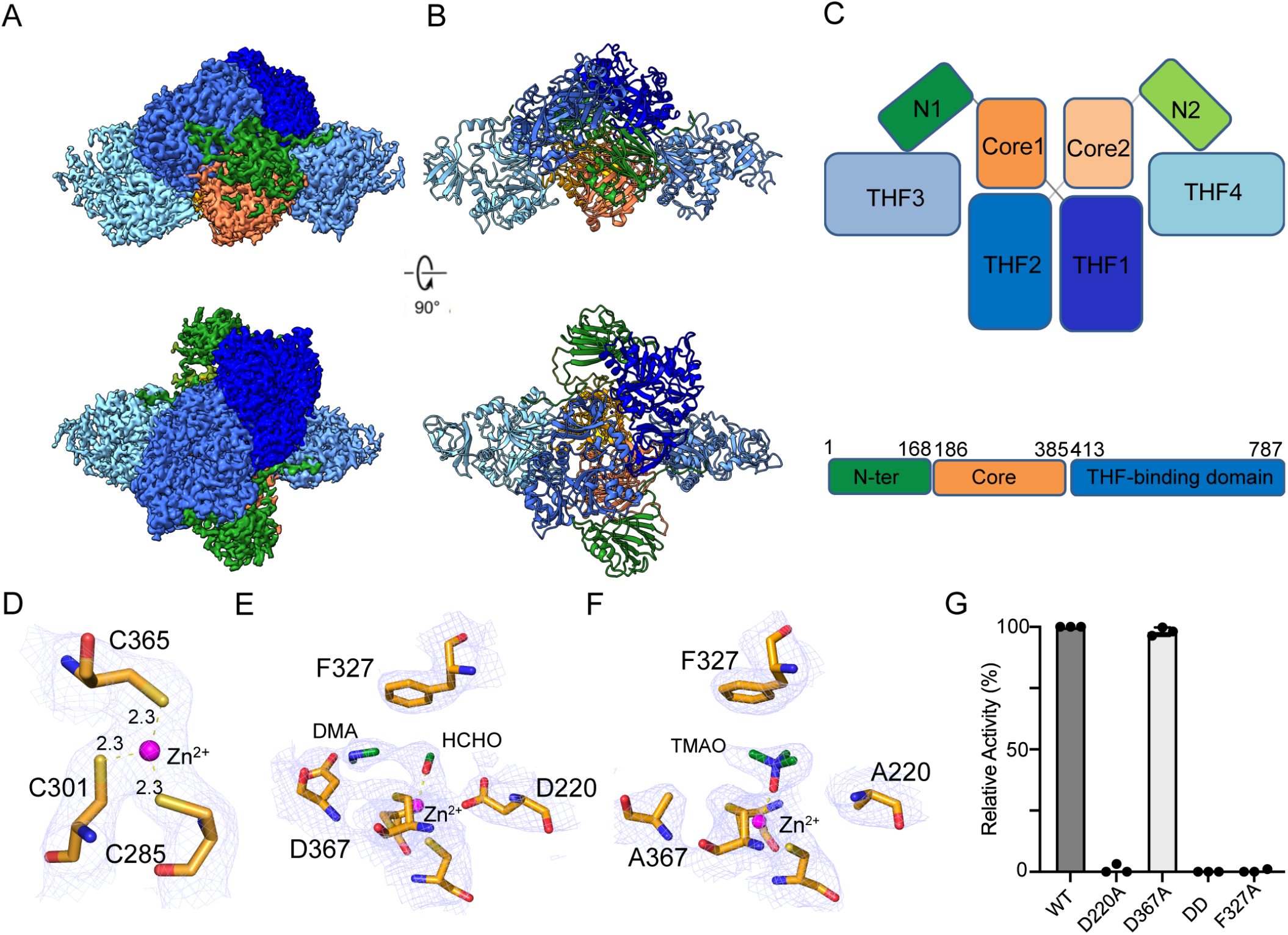
Overall architecture of TDM determined by cryo-electron microscopy. (A) Representative cryo-electron microscopy (EM) maps. (B) Overall structure of the complex, with each subunit colored differently. TDM adopts a 2 + 2^1/2^ complexation. The two half-domains of the complex are the C-terminal domains. (C) (C) Schematic illustration of TDM complexation. TDM monomers with N-terminal, Core, and C-terminal domains are colored forest, orange, and blue, respectively. The same coloring is used in panels A and B. (D) Close-up view of the 3Cys:Zn^2+^ binding motif. (E) Close-up view of the interactions between DMA and HCHO products with the 3Cys:Zn^2+^ motif. (F) The TMAO substrate is trapped in the 3Cys:Zn^2+^ active site of the double D220A/D367A mutant. The density maps are shown and contoured at 3.0 σ with a carve of 2 Å. Bound small molecules are shown as ball-and-stick representations, amino acid residues as sticks, and hydrogen bonds as dashed lines. (G) Relative activity of TDM variants.

Each catalytic core domain contains a conserved 3Cys:Zn^2+^ motif (C285, C301, C365) essential for protein stability (Figure 2D, S4). Substitution of any of these cysteines with alanine led to aggregation, underscoring their structural importance. Elemental analysis (EDS and ICP–MS) confirmed zinc incorporation but excluded iron, yielding a Zn: protein ratio of ∼1:2 (Figure S5A, B). This motif is analogous to Zn^2+^ centers in cytidine deaminases,^[10,11]^ alcohol dehydrogenases, and other Zn metalloenzymes.^[12,13]^

### Zn^2+^-Dependent Organization at the Core Catalytic Domain

In TDM, Zn^2+^ occupies a single, well-defined site within the core catalytic domain and plays a central non-redox role in substrate binding and organization of the active-site environment. Cryo-EM analysis of the substrate-bound TDM complex reveals TMAO positioned within the Zn^2+^-containing active site through coordination of its oxygen atom with the metal center, while the surrounding conserved residues D220, F327, and D367 contribute to substrate recognition and stabilization (Figure 2D-F). These interactions define the substrate-bound state of the catalytic pocket and are summarized in Scheme 1, which illustrates the experimentally observed interactions between TMAO and the Zn^2+^-containing active site.

The cryo-EM structure of the TDM:DMA:HCHO complex shows both products retained within the active site, whereas in the D220A/D367A:TMAO mutant structure, the substrate remains trapped without substantial conformational changes relative to the wild-type enzyme (Figure 2E, F). Consistent with these observations, the F327A and D220A variants exhibit markedly reduced enzymatic activity, underscoring their importance in substrate recognition and active-site organization (Figure 2G). In contrast, mutation of D367, which participates in hydrogen bonding with the DMA product, results in a more modest (∼10%) reduction in activity, consistent with a role in product stabilization rather than substrate binding or activation (Figure 2E–G). Together with the substrate-bound structure, these product-associated interactions complete the structural framework summarized in Scheme 1, illustrating how the Zn^2+^-containing active site accommodates both the substrate and the reaction products during TMAO turnover. While the detailed chemical mechanism remains to be established, these structures provide a framework for understanding substrate recognition, product stabilization, and active-site organization in TDM.

**Scheme 1.**
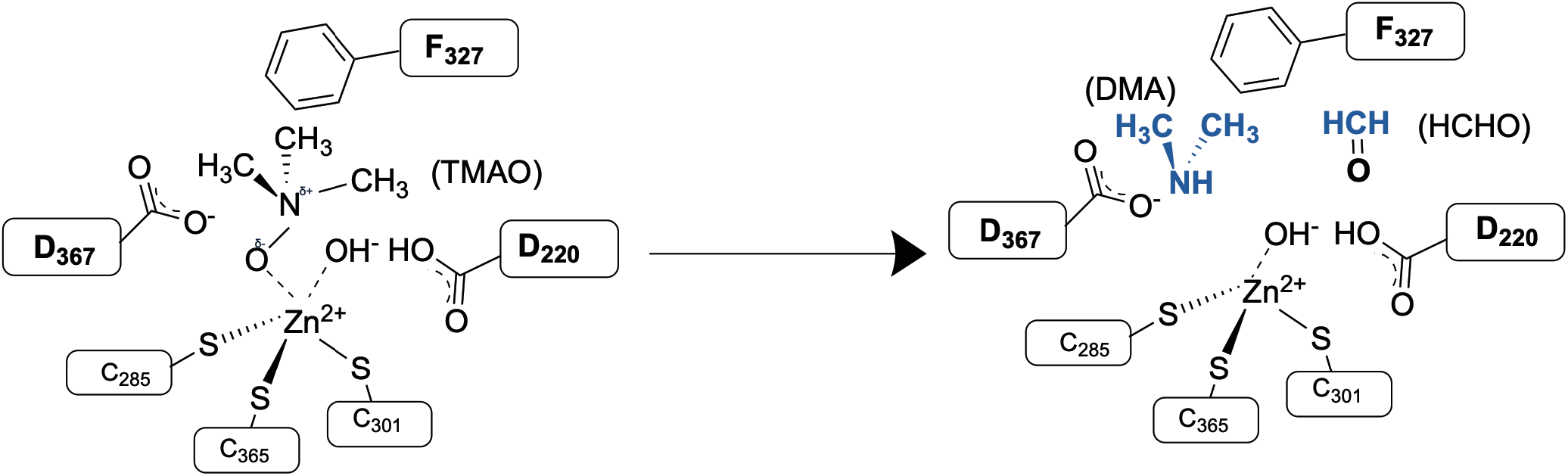
Interactions of the substrate (TMAO) and reaction products within the Zn^2+^-containing active site of TDM. TMAO is coordinated within the active site through interactions with Zn^2+^ and surrounding conserved residues. The proposed product-binding states illustrate interactions of dimethylamine (DMA) and formaldehyde (HCHO) with residues lining the active-site pocket following turnover.

Notably, we found no structural or biochemical evidence for an additional iron-binding site in TDM. Previous studies of TDM from *Methylocella silvestris* proposed the involvement of both Zn^2+^ and non-heme Fe^2+^ based on metal-content analyses and mutagenesis experiments; however, in the absence of experimentally determined structural information, the locations and coordination environments of these metals could not be directly resolved.^[7,14]^ In contrast, our ICP-MS and cryo-EM analyses consistently identified a single Zn^2+^-binding site within the core catalytic domain. Furthermore, expression of TDM in LB medium supplemented with 0.5 mM Fe(NH4)2SO4 (Figures S6A and S6B) did not reveal detectable Fe incorporation or the presence of an additional metal-binding site. Collectively, these data support a single Zn^2+^-containing active site in our TDM preparation, and suggest that its metal composition differs from that proposed for the *M. silvestris* enzyme. While Zn^2+^ is structurally associated with substrate binding and organization of the active-site environment, the detailed chemical basis of TMAO demethylation remains to be established.

### Architecture and Binding Properties at THF-Binding Domain

The C-terminal THF-binding domain is positioned on the intramolecular face of the TDM complex, and forms extensive electrostatic interactions that are essential for stabilizing the central dimer interface (Figure S7A, B).^[15,16]^ Structural comparison revealed that this domain adopts a conserved folate-binding fold closely resembling those of previously characterized THF-dependent proteins. Superposition with the T protein of the glycine cleavage system (PDB ID: 1WOO) yielded an RMSD of 1.18 Å over 371 Cα atoms (Figure S7C), indicating a high degree of structural conservation despite limited sequence identity.^[15]^ In addition, the overall architecture is consistent with the folate-binding domain of formiminotransferase-cyclodeaminase, another THF-dependent enzyme that utilizes a dedicated folate-binding module and has been proposed to facilitate intermediate transfer between catalytic domains.^[16]^

Cryo-EM density at the folate-binding site was consistent with that of THF binding, although resolution limitations and partial occupancy prevented complete ligand assignment (Figure 3A). Based on the structural homology, we modeled THF in the central cavity of the ring structure (Figure 3B). The pterin moiety of THF projected into the interior cavity, forming hydrophobic contacts with the aromatic folate-binding pocket (Figure 3B). Two positively charged residues, R480 and K782, located near the entrance of the binding funnel, likely engaged the polyglutamate tail via hydrogen bonding, complemented by π-stacking from F582 and F672 (Figure 3B). Similar binding modes have been observed in enzymes, such as T-proteins, aminotransferase domains, and dimethylglycine dehydrogenase.^[17,18,19]^ ITC binding isotherms show a clear interaction between THF and TDM, with negligible heat change in the buffer control, confirming binding specificity. The dissociation constant (*Kd*) was determined to be approximately 149 nM (Figure 3C, S8).

**Figure 3.**
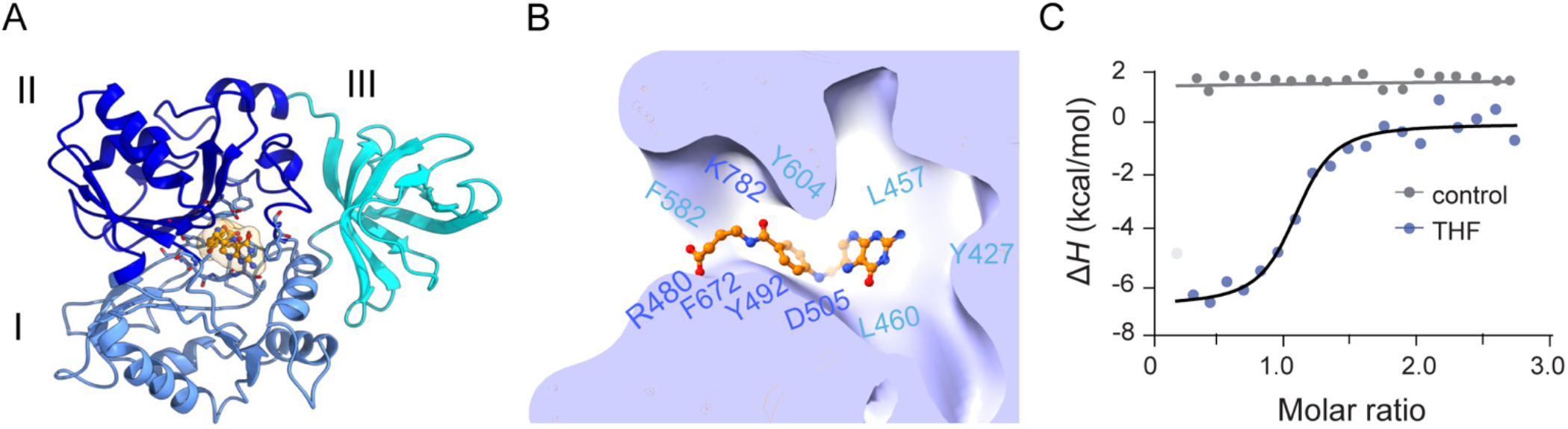
THF binding to TDM. (A) The THF-binding domain forms a cloverleaf of three subdomains (I-III), creating a central cavity that accommodates THF. EM density for THF is shown as a surface. (B) Close-up of the binding pocket highlighting hydrophobic and hydrogen-bonding residues that stabilize THF, colored according to their subdomain location in panel A.(C) ITC analysis confirms specific THF binding to TDM (blue), with negligible signal in buffer control (gray).

### A Substrate Channel for Formaldehyde Transfer

High-resolution cavity mapping of the TDM:DMA:HCHO: THF complex revealed a continuous intramolecular tunnel that connects the Zn^2+^-containing demethylase pocket to the distal THF-binding site (Figure 4A). This conduit begins at the formaldehyde release position within the catalytic pocket at the core domain (HA, HB) and terminates near the folate pterin ring at the C-terminal domain (TA, TB). The channel is predominantly lined with acidic residues and polar side chains, generating a strongly negative electrostatic potential (Figure 4A). Such electrostatics likely stabilize the partial negative oxygen of HCHO, guiding it through the tunnel while excluding bulk anions and minimizing leakage.

**Figure 4.**
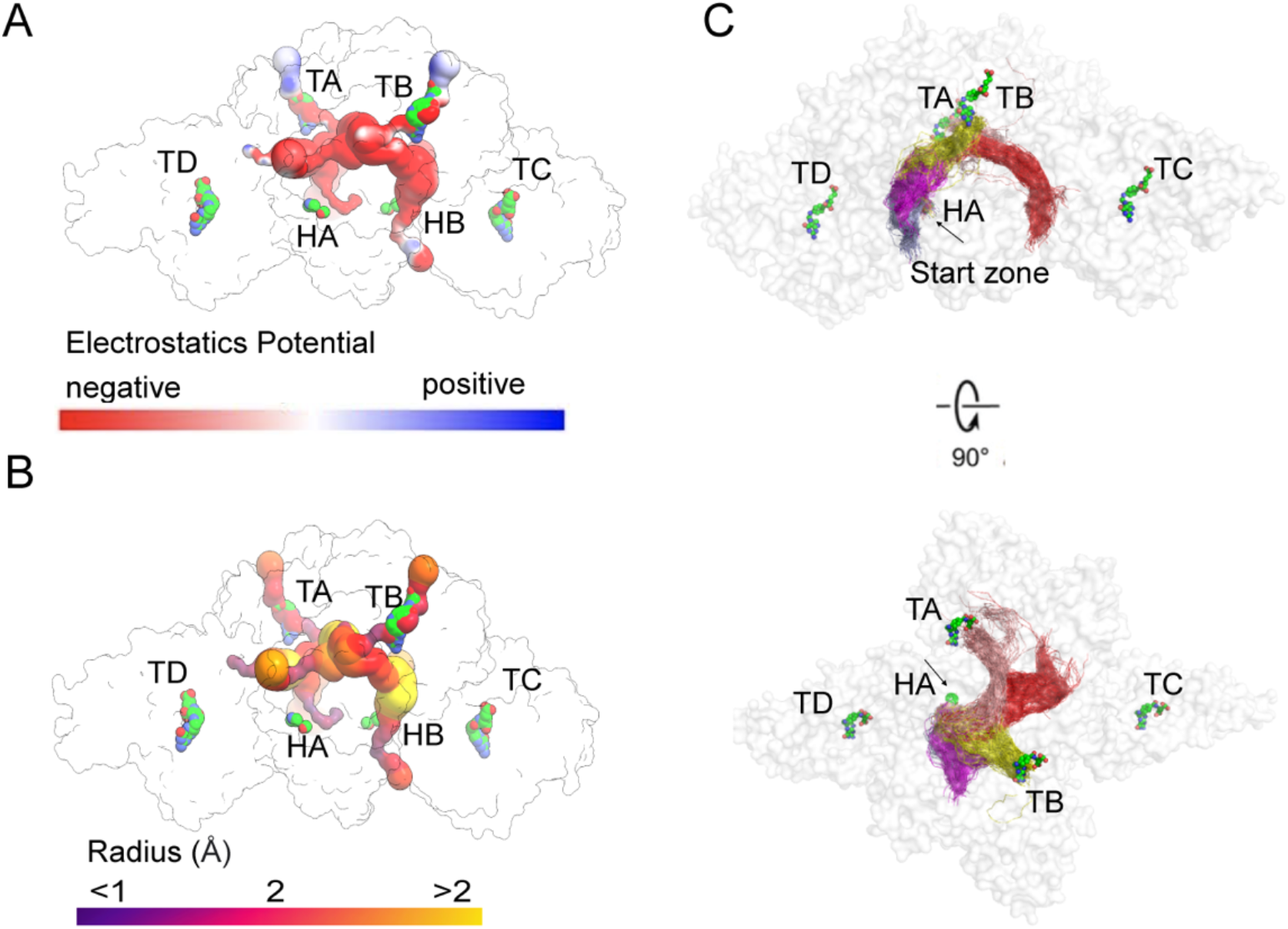
HCHO channels from the active site to the THF-binding site in TDM complex. (A, B) Surface representations of the channels within the TDM complex colored according to either electrostatic potential or radius. (C) Tunnel analysis of a CG MD simulation suggests that HCHO in each chain can access the THF binding sites in chains A and B. The five top-ranked paths for chain A are shown in different colors. THF, DMA, and HCHO are depicted as spheres colored by the atom type. TA, TB, TC, and TD: THF molecules in chains A, B, C, and D, respectively; HA: chain A, HCHO.

Geometrically, the tunnel exhibits a radius ranging between 2.6 and 5.0 Å, with the narrowest bottleneck of 2.6 Å positioned just before the folate-binding cavity (Figure 4B). This matches closely with the van der Waals dimensions of formaldehyde (∼2.5 Å), suggesting evolutionary fine-tuning for selective transfer of this reactive one-carbon intermediate. Importantly, no alternative solvent-accessible exits were detected, indicating that HCHO is obligatorily channeled to the THF site rather than diffusing freely.

Coarse-grained target molecular dynamics simulations further supported this interpretation. Independent trajectories consistently identified two persistent tunnels, each connecting the Zn^2+^ demethylation sites (HA, HB) to the corresponding folate-binding sites (TA, TB) in the two protomers (Figure 4C, S9). Within these trajectories, formaldehyde molecules entered the channel from the catalytic site, transiently interacted with acidic residues and ordered water molecules, and passed through narrow gating residues that rearranged to allow transit (Figure 4C). This dynamic yet persistent tunnel suggests a gated channeling mechanism that balances selectivity with throughput.

The channeling mechanism in TDM bears resemblance to classical substrate-channeling systems but with distinct architectural solutions. In tryptophan synthase, an ∼25 Å hydrophobic tunnel delivers indole between two active sites,^[20,21]^ whereas the glycine cleavage system employs a swinging lipoate arm on the H-protein to transfer intermediates between catalytic partners.^[22,23]^ Dimethylglycine dehydrogenase also channels formaldehyde to THF, but does so through surface-proximal transfer rather than via a buried conduit,^[9,19]^ while the PaaZ enzyme in phenylacetate catabolism uses electrostatic pivoting, in which a domain-swapped CoA arm swings the intermediate between domains, guided by conserved positive residues.^[22]^ By contrast, TDM relies on a fully enclosed, negatively charged tunnel, a unique design optimized for formaldehyde, a small, highly diffusible, and cytotoxic intermediate.

### Role of the N-terminal domain in TDM assembly and function

The function of the N-terminal domain in TDM has remained unclear, and sequence alignment across homologs reveals low evolutionary conservation in this region (Figure 5A). Cryo-EM analysis revealed two distinct interaction modes between the N-terminal and C-terminal domains, including an unexpected interdomain interface between the N-terminal region and a peripheral THF-binding domain (Figure 5B, S10, Video S1). This interface appears to tether the flexible N-terminus to the THF-binding domain, forming a structural bridge that may regulate the conformational transitions required for complex assembly and efficient substrate transfer. The interaction is stabilized by aromatic stacking between P14 and W476, charge interaction, along with extensive hydrophobic contacts that promote proper domain packing and interdomain communication (Figure 5B, S9).

**Figure 5.**
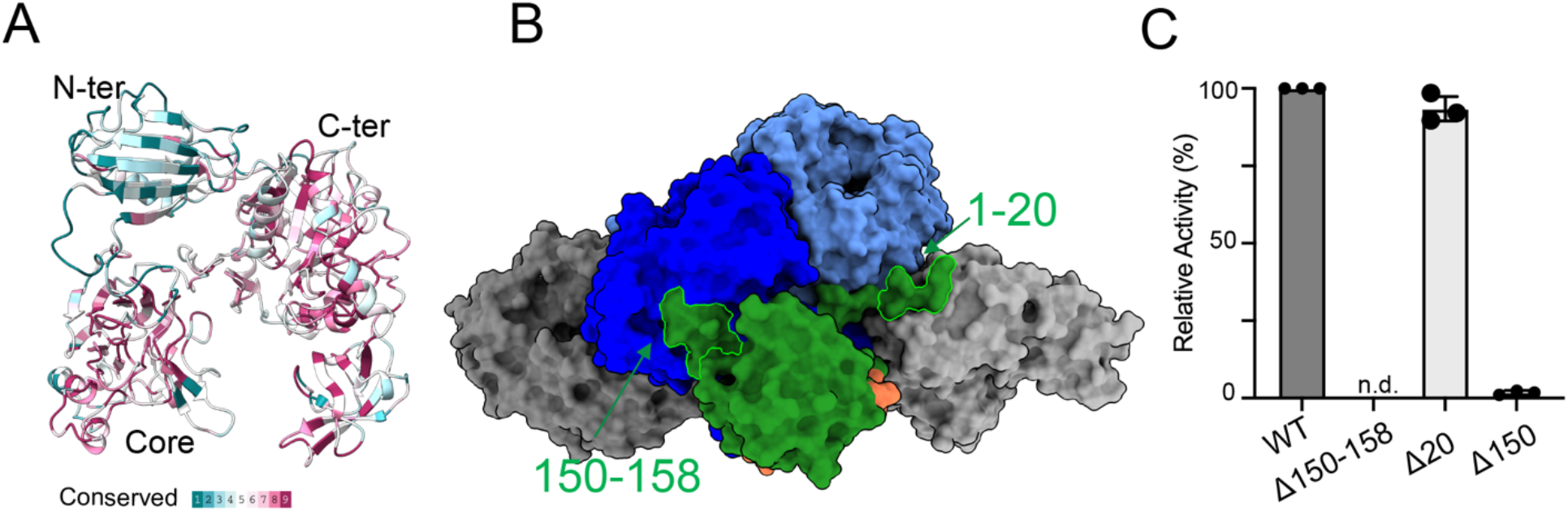
The N-terminal domain is involved in the structural stability and assembly of proteins. **(A)** The evolutionary conservation of residue positions on the TDM structure is estimated based on phylogenetic connections among homologous sequences. (B) N-terminal domain was depicted as forest. Residues ranging from 1-20, 150-158 were highlighted. (C) Relative TDM activity of TDM mutants compared to that of WT variant.

Functional analyses further support this structural coupling. Deletion of the entire N-terminal region (Δ1–150) or the interfacial segment (Δ151–158) led to severe aggregation and loss of soluble protein, while removal of only the first 20 residues (Δ1–20) caused a modest (∼10%) reduction in catalytic turnover (Figure 5C). These results indicate that the N-terminal domain not only stabilizes the quaternary architecture but may also facilitate allosteric coupling between the catalytic and folate-binding domains. Such dynamic coupling likely ensures efficient handoff of formaldehyde through the internal tunnel, synchronizing demethylation and one-carbon transfer reactions within the bifunctional TDM complex.

## CONCLUSION

Our results establish TDM as both a bifunctional enzyme and an integrated substrate-channeling system. By coupling TMAO demethylation with THF-dependent one-carbon transfer, TDM not only produces DMA and methylene-THF but also ensures immediate capture of formaldehyde, a highly reactive and cytotoxic intermediate. Structural, biochemical, and computational analyses reveal that this is achieved through a fully enclosed, negatively charged conduit that links the Zn^2+^ catalytic core to the folate-binding site, thereby enhancing catalytic efficiency and preventing metabolite leakage.

Beyond defining the molecular basis of TDM activity, these findings highlight a general design principle: substrate channeling can evolve distinct architectures, hydrophobic tunnels, swinging arms, or electrostatic conduits, tailored to the physicochemical properties of the intermediate. The TDM structure expands this paradigm by revealing a tunnel optimized for formaldehyde transfer, underscoring microbial strategies for detoxification and metabolic efficiency. More broadly, this work provides a framework for re-engineering channeling systems in biocatalysis, synthetic biology, and microbiome modulation, where controlling the fate of reactive intermediates is central to both metabolic innovation and therapeutic applications.

## EXPERIMENTAL DETAILS

### TDM Construction, Expression, and Purification

The gene encoding full-length TDM (1-787) (WP_26356686) from *Paracoccus* sp. DMF was optimized for *E. coli* codons and cloned into the pET21a(+) plasmid with either a C-terminal 6xHis tag or an N-terminal strepII tag using NdeI and XhoI restriction enzymes. Site-directed mutagenesis involving substitutions or deletions was performed using specific PCR primers (Universal, USA). Plasmid isolation and purification from E. coli were performed using a DNA spin kit (Qiagen, Germany), and the identity of the DNA constructs was confirmed by sequencing (Genewiz, USA).

The TDM protein variants were expressed in *E. coli* BL21 (DE3) cells cultured in LB medium at 37 °C. Induction was achieved with 0.25 mM isopropyl β-D-1-thiogalactopyranoside (IPTG) at 18 °C for 16 h. Protein purification was performed by affinity chromatography using either Ni-NTA resin (Qiagen, Germany) or a strep-tag column (Cytiva, USA) in buffer A (20 mM Tris-HCl, pH 7.5, 250 mM NaCl, 1 mM TCEP, 300 mM imidazole, and 5% glycerol). Further purification steps included passage through a 2 × 5 mL HiTrap Q column (GE Healthcare, USA) using buffer B (20 mM Tris-HCl pH 7.5, 200 mM NaCl, 1 mM TCEP, and 5% glycerol) for equilibration, and buffer C (20 mM Tris-HCl pH 7.5, 1000 mM NaCl, 1 mM TCEP, and 5% glycerol) for elution. Final purification was achieved using a Superdex 200 increase 10/300 GL column (Cytiva, USA) in buffer D (20 mM Tris-HCl, pH 7.5, 150 mM NaCl, and 5% glycerol). The purity of the relevant fractions was assessed by SDS-PAGE.

### TDM Activity Assay

Enzyme activity was assessed by measuring HCHO production from TMAO breakdown as described previously with minor modifications.^[23]^ Briefly, enzymatic activity was evaluated in multiple buffering systems spanning a range of pH values. Maximal activity was observed in 10 mM MES buffer (pH 6.0) supplemented with 150 mM NaCl; this buffer was subsequently used for all kinetic analyses. Standard activity assays were performed in triplicate at room temperature. Each 50 μL reaction contained 2.5 μg of purified TDM in MES buffer and was initiated by the addition of TMAO to a final concentration of 10 mM. Reactions were incubated for 10 min, a time point confirmed to be within the linear range based on preliminary time-course experiments. For HCHO detection, 10 μL of the reaction mixture was combined with 25 μL of freshly prepared 0.2% (w/v) Purpald reagent dissolved in 1 M NaOH and 215 μL of Milli-Q water in a 96-well plate. After incubation for 20 min at room temperature, absorbance at 540 nm was measured using a SpectraMax G5 microplate reader. Formaldehyde concentrations were calculated from a standard curve generated using analytical-grade formaldehyde (0–180 μM). The Michaelis–Menten constant (Km) and maximum velocity (Vmax) were determined using the PRISM v.10.0.0 software. To evaluate HCHO reduction in the presence of THF, reactions were conducted with 5 mM TMAO and varying concentrations of THF ranging from 0 to 150 µM.

For the analysis of DMA, CH2-THF products, and other compounds, high-performance liquid chromatography (HPLC) coupled with Mass Spectrometry (MS) was employed. The analysis was performed using an Agilent 1290 Infinity II LC system coupled with an Agilent 6470 series QQQ mass spectrometer. Waters Acquity UPLC with a bridged ethylene hybrid hydrophobic interaction chromatography (2.1 mm × 150 mm, 1.7 µm) column was used for separation. The mobile phases were as follows: (A) 90% acetonitrile, 5% isopropanol, and 5% 200 mM ammonium formate (pH 3) and (B) 90% water, 5% acetonitrile, and 5% 200 mM ammonium formate (pH 3). The LC gradient was: 0 min, 0% B; 2 min, 0% B; 7 min, 98% B; 7.5 minutes, 98% B; 8 min, 0% B; 15 min, 0% B, with a flow rate of 0.3 mL/min.

Mass spectrometric analysis was performed using multiple reaction monitoring (MRM) in the positive electrospray ionization (ESI) mode. The ESI interface settings were as follows: gas temperature, 325°C; gas flow rate, 7 L/min; nebulizer pressure, 45 psi; sheath gas temperature, 250°C; sheath gas flow rate, 7 L/min; capillary voltage, 3800 V; nozzle voltage, 1000 V; and ΔEMV voltage, 300 V. The concentrations of TMAO, DMA, THF, and CH2-THF were determined using internal deuterated standards (TMAO-d9 and DMA-d6), and the data were analyzed using the Agilent MassHunter Quantitative Analysis (Version B.08.00).

### Cryo-EM data processing and structure determination

Image processing and structure determination involve several steps. Motion correction and image summation were conducted using MotionCor,^[24]^ and defocus values were estimated using Gctf.^[25]^ Particle picking was performed using the DoG Picker,^[26]^ followed by initial reference-free 2D classification in CryoSPARC v.4.2.0.^[27]^ Representative 2D class averages were selected, and particle refinement was performed through multiple rounds of 2D classification. Subsequently, an initial ab initio reconstruction was performed. The particles underwent 3D classification into five classes using a low-pass filtered initial reconstruction at 20 Å as a reference model. Multiple rounds of 3D classification were performed, and the optimal subset displaying clear structural features underwent heterogeneous refinement over six rounds, resulting in a high-quality, refined subset. These refined particles underwent homologous refinement, yielding a map with global resolution indicated by a Fourier shell correlation (FSC) of 0.143. The final volume map was computed with C2 symmetry using CryoSPARC, and local resolution was assessed using Phenix. The initial apo model of the TDM was built in Coot v0.8.9,^[28]^ referencing the alphafold structure.^[29]^ Monomeric TDM structures were fitted into cryo-EM electron density maps using UCSF ChimeraX,^[30]^ with subsequent iterative manual adjustments and rebuilding in Coot, including modeling of the N-terminal domain. Manual refinement was based on the electron density map quality in Coot, followed by real-space refinement in Phenix.^[31]^ Validation included the analysis of FSC curves to assess model-map agreement and the evaluation of atomic model geometry using MolProbity.^[32]^ Detailed refinement statistics are provided in Table 1. The pore-lining surfaces and receptor channels were analyzed using the MOLE software.^[33]^ Visual representations were created using UCSF ChimeraX and PyMOL.^[34]^

### Molecular Dynamics simulations of enzyme tunnelling

Coarse-grained (CG) MD simulations of TDM were conducted using the GROMACS 2021.5 and Martini3 force fields with the Gō model.^[35]^ The CG coordinates were generated from the TDM:DMA: HCHO structure via martinize2,^[36]^ with the secondary structure assigned using DSSP.^[37]^ The Gō contact map was created using the GōContactMap server with a Cα–Cα distance cutoff of 0.3–1.1 nm and Gō potential of 9.414 kJ·mol^−1^.^[35]^

The simulations employed periodic boundary conditions, with short-range interaction cutoff at 1.1 nm and long-range electrostatics handled by the reaction-field method.^[38]^ The system was maintained at 316.16 K and a pressure of 1 bar. Initial energy minimization was performed until the forces were below 10 kJ·mol^−1^·nm^−1^, followed by solvation using INSANE^[39]^ and addition of 0.15 M NaCl. The system contained 1324 Na^+^/Cl^−^ beads and 110,246 water beads in the simulation box.

The system was equilibrated for 5 ns with position restraints on the backbone beads, followed by 10 ns of unrestrained equilibration. The 10 µs production run used the velocity-rescale thermostat, and Parrinello-Rahman barostat, with a 20 fs time step and coordinates saved every 500 ps for the production run. ^[40,41]^ Tunnel calculations were performed using CAVER3.0.^[42]^ The starting points for the tunnel calculations were defined by residues near HCHO (A282, T369, and S378). A 2.5 Å probe radius and time sparsity of 1 were applied to avoid redundant tunnel generation. The detected tunnels were clustered using a threshold of 5.0 and a frame reweighting coefficient of 1. Protein stability was evaluated by calculating RMSD, RMSF, and Rg using *gmx rms, gmx rmsf*, and *gmx gyrates*, respectively.

### Isothermal Calorimetry (ITC)

ITC measurements were performed using a NANO ITC system (TA Instruments) to evaluate the binding affinities of TDM and THF at 20 °C. The compound solution was equilibrated with protein buffer D, and all protein and compound solutions were prepared in the same degassed buffer before use. ITC cells were loaded with 250 μl of TDM protein (5 μM) and titrated with THF (80 μM). Titration involved injecting 1 μl of the substrate into the protein solution, followed by 19 injections of 2.45 μl each. The stirring rate was set at 200 rpm. For control experiments, the buffer was injected into the protein alone. Equilibrium association constants were calculated by fitting reference-corrected data using the binding model supplied by the manufacturer.

### Energy-dispersive X-ray spectroscopy analysis (EDAX)

Ten microliters of TDM protein at a concentration of 100 µM was dropped on the silicon wafer and vacuum dried. After drying, the samples were gold-coated (∼ 10 nm) and analyzed using a CARL ZEISS EVO 50. Buffer was used as a control experiment.

### Inductively coupled plasma-mass spectrometry (ICP-MS)

ICP-MS experiments were conducted using trace metal-grade nitric acid (HNO3, 3% v/v), which was purified in-house by sub-boiling point distillation, as the sample matrix. ICP-MS analyses were performed using an Agilent Technologies 7500 ICP-MS instrument, while ICP-OES was conducted using a Perkin Elmer Optima 5300DV Optical Emission Spectrometer. Protein samples at approximately 1.0 mg/mL, together with buffer-only controls, were initially screened by ICP–MS using a stitched wide-window survey scan to approximate a full metal mass scan for the detection of metal-associated isotopes, while also identifying predominant elemental species and potential spectral interferences. Calibration standards were freshly prepared by diluting the stock solutions (Sigma-Aldrich) in 3% HNO3 and doubly deionized water. Approximately 2.4 mg of protein was diluted in a 3% HNO3 matrix for metal analysis, and zinc concentrations were measured using the emission line at 213.857 nm.

## Author Contributions

TT, KD, and SM conducted cellular and biochemical studies. YH conducted HPLC-MS. SQ and JR performed MD simulations. TT, GR, and RS were involved in designing the experiments and developing this project. TT and RS were involved in the structure determination, map interpretation, and result analysis. All authors participated in manuscript preparation and review.

## Acknowledgements

Special thanks go to Dr. Robert Stahelin for granting access to the ITC machine; Dr. Na Gou for ICP-MS analysis; Dr. T. Klose and Dr. F. Vago for cryo-EM, and S. Wilson for computation. We acknowledge the Purdue cryo-EM facility for access to instrumentation for data collection, and the New Zealand eScience Infrastructure for high-performance computing.

## Table of Contents

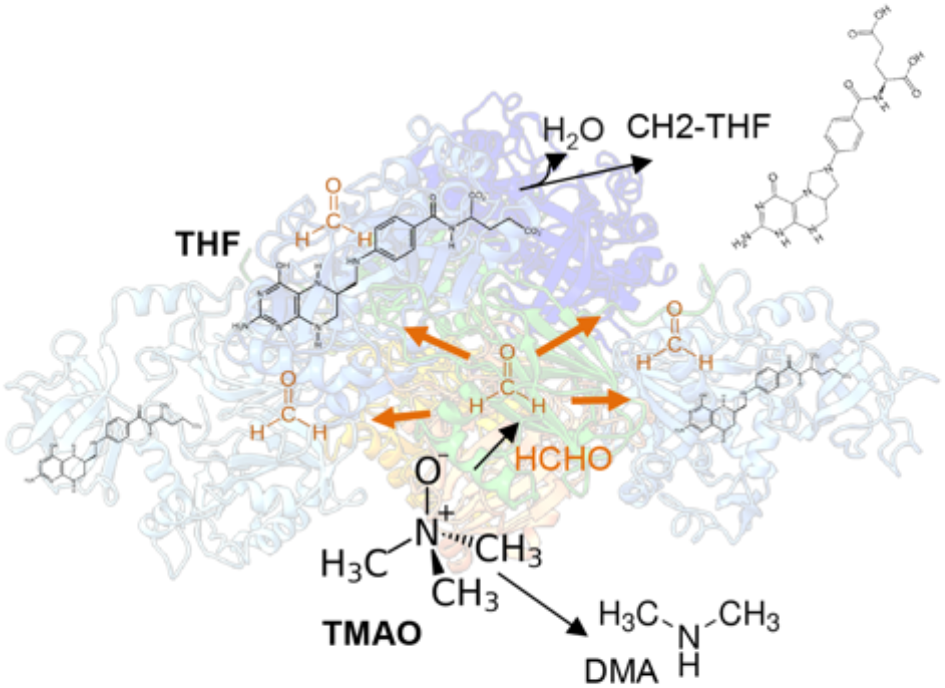

