## supplementary for "Bifunctional Architecture Enables Substrate Catalysis and Channeling in *Paracoccus* TMAO Demethylase" (from another version of this preprint: version 2, 2026-02-18)

Supplementary FigureS

- Figure S1. Purification and biochemical characterization of TDM
- Figure S2. The cryo-EM data processing pipeline for TDM structures carried out using cryoSPARC
- Figure S3. Dimerization in the core domains
- Figure S4. Sequence alignment of TDM from Paracoccus sp. and other organisms.
- Figure S5. Zinc association with the TDM protein
- Figure S6. paraTDM lacks an iron-binding site
- Figure S7. THF-binding domain of TDM
- Figure S8. The binding of THF to TDM was determined by ITC
- Figure S9. Molecular Dynamics simulation
- Figure S10. Two distinct binding modes between the N-terminal domain and the THF-binding domains

Supplementary Video

- N-terminal domain facilitates recruitment of the peripheral THF-binding domain

**SUPPLEMENTARY FIGURES**
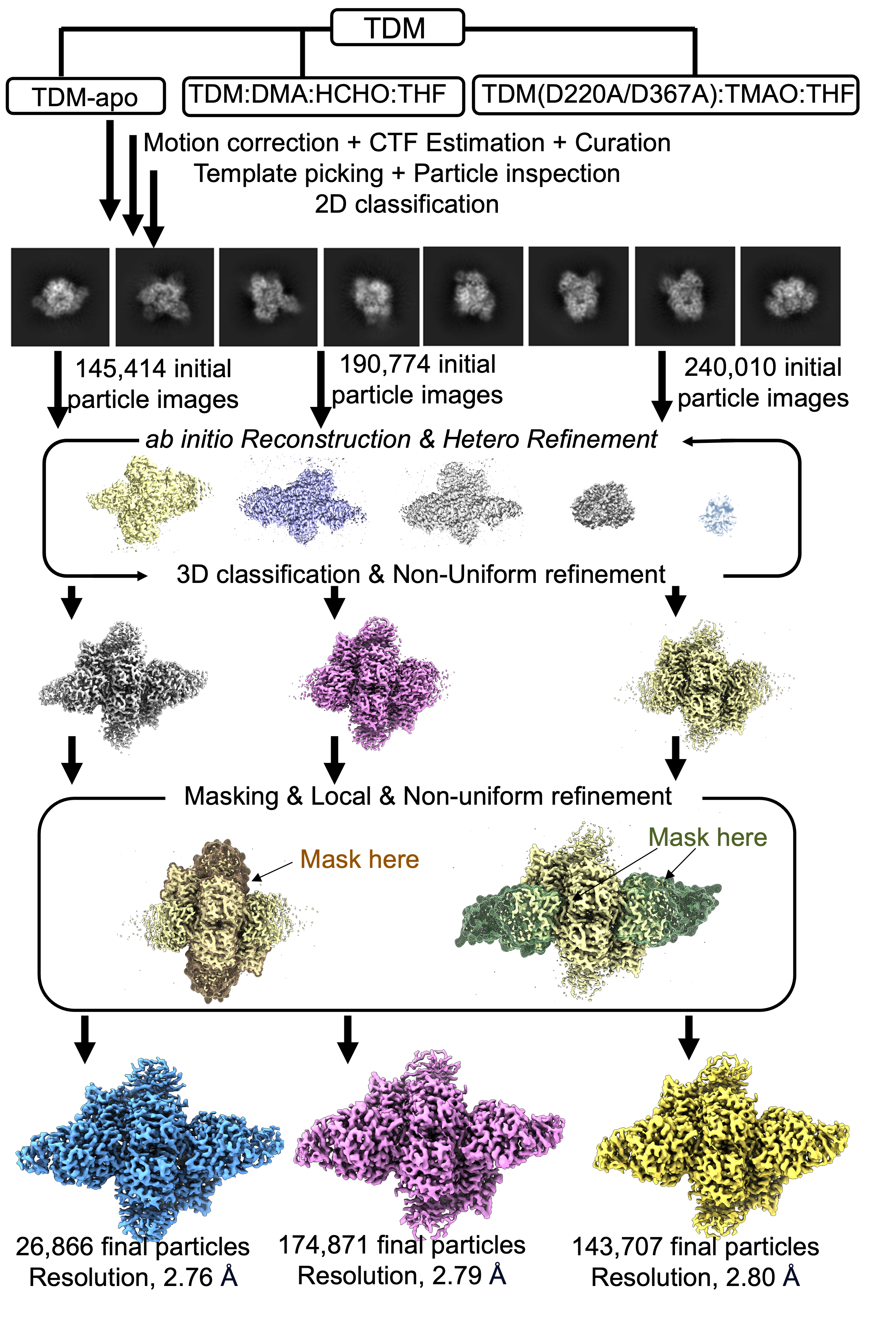

Figure S1. Purification and biochemical characterization of TDM. (A) Protein samples were fractionated using a Superdex 200 10/300 column at varying salt concentrations. The peaks below 10 mL were in the void volume and aggregated. At 150mM NaCl, a peak around 12.5 ml shows the best activity. (B) SDS-PAGE was used to assess sample purity. The gel shows the band under the gray and black fractions (corresponding to elution volumes marked in A) at 150mM NaCl. Note that all peaks have TDM. The arrow on the right shows a band corresponding to the molecular weight of TDM. (C) TDM catalytic activity at different pH values (20 mM acetate buffer for pH of 4.0-5.0; 20 mM MES buffer for pH of 5.5-6.5; 20 mM Tris-HCl buffer for pH of 7.0-9.0) at 37 °C for 30 min and (D) Temperature conditions in a buffer of 20 mM MES pH 6.5, 150 mM NaCl. Relative activity was measured in relation to the maximum activity, which was set at 100%. All kinetic experiments were performed in triplicates.

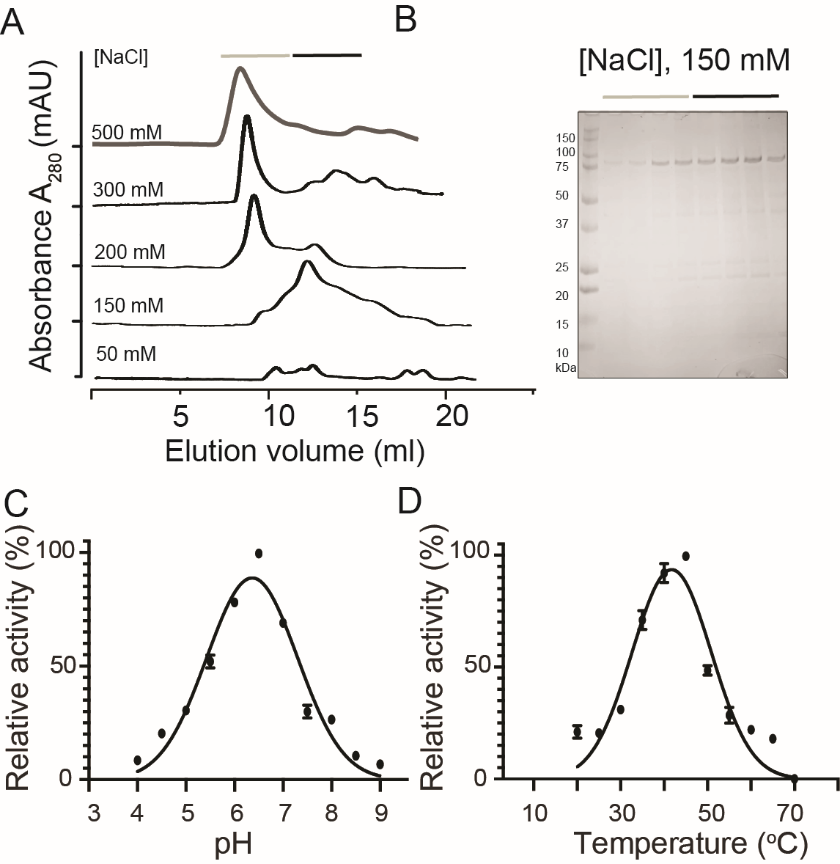

**Figure S2. The cryo-EM data processing pipeline for TDM srtuctures carried out using cryoSPARC.** Following iterative rounds of particle selection, classification, and refinement, a final subset comprising 10-60% of the initially exported particles was used to reconstruct the 3D EM map. The resulting map was subjected to local resolution estimation using the PHENIX, which revealed a resolution range spanning from 2.0 to 5.0 Å across different regions of the structure. Both the global and local resolution maps were subsequently visualized and analyzed using UCSF ChimeraX to facilitate model interpretation and validation.

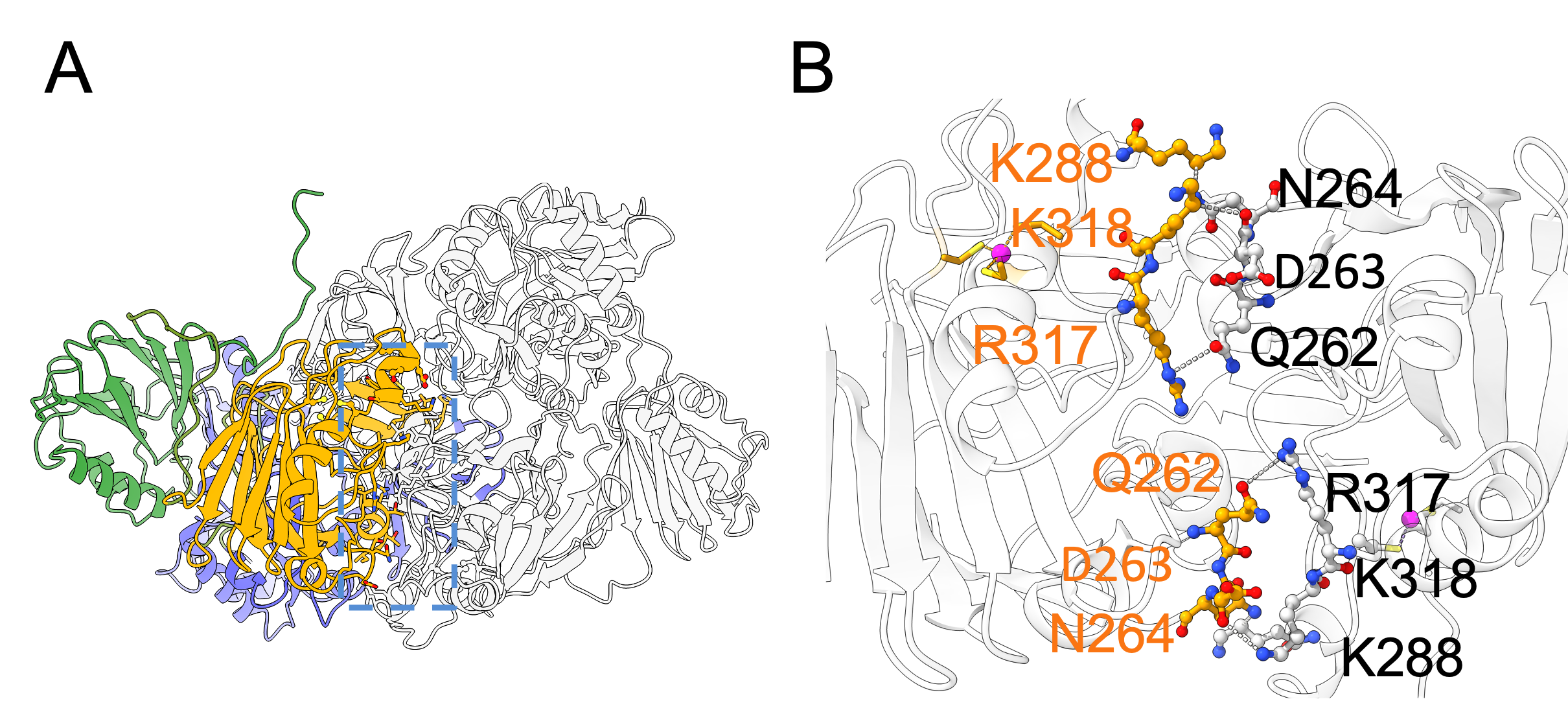

Figure S3. ****Dimerization of the core domains in the TDM complex.**** (A) The central dimer interface is mediated by residues located within the core domain of each TDM monomer. For clarity, the N-terminal, core, and C-terminal domains are shown in forest green, light orange, and light blue, respectively. This structural arrangement highlights the spatial organization of the domains contributing to dimer formation. (B) The interface between the two core domains is stabilized through a network of interactions, including hydrogen bonds and hydrophobic contacts. Key residues involved in these interactions are positioned symmetrically across the interface, contributing to the overall stability and specificity of the dimeric assembly.

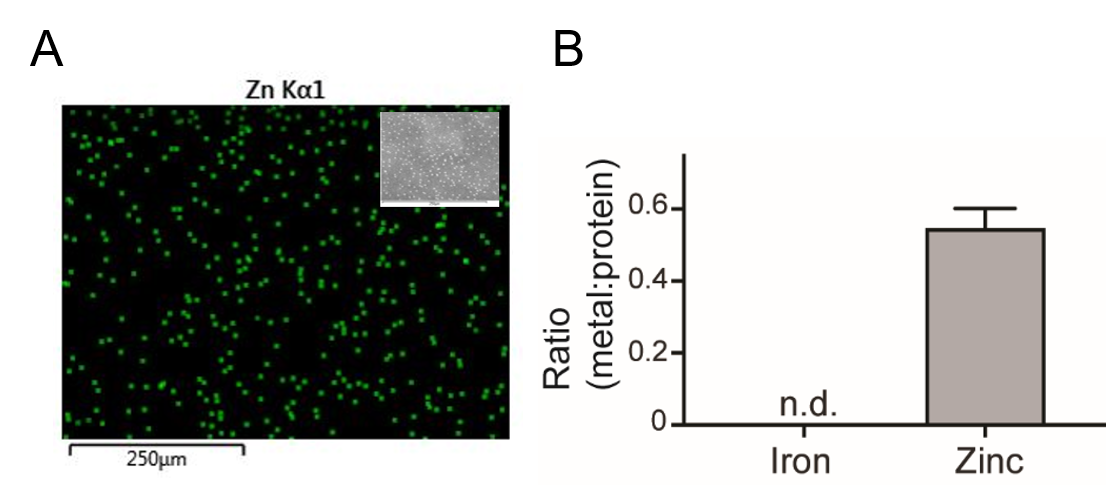

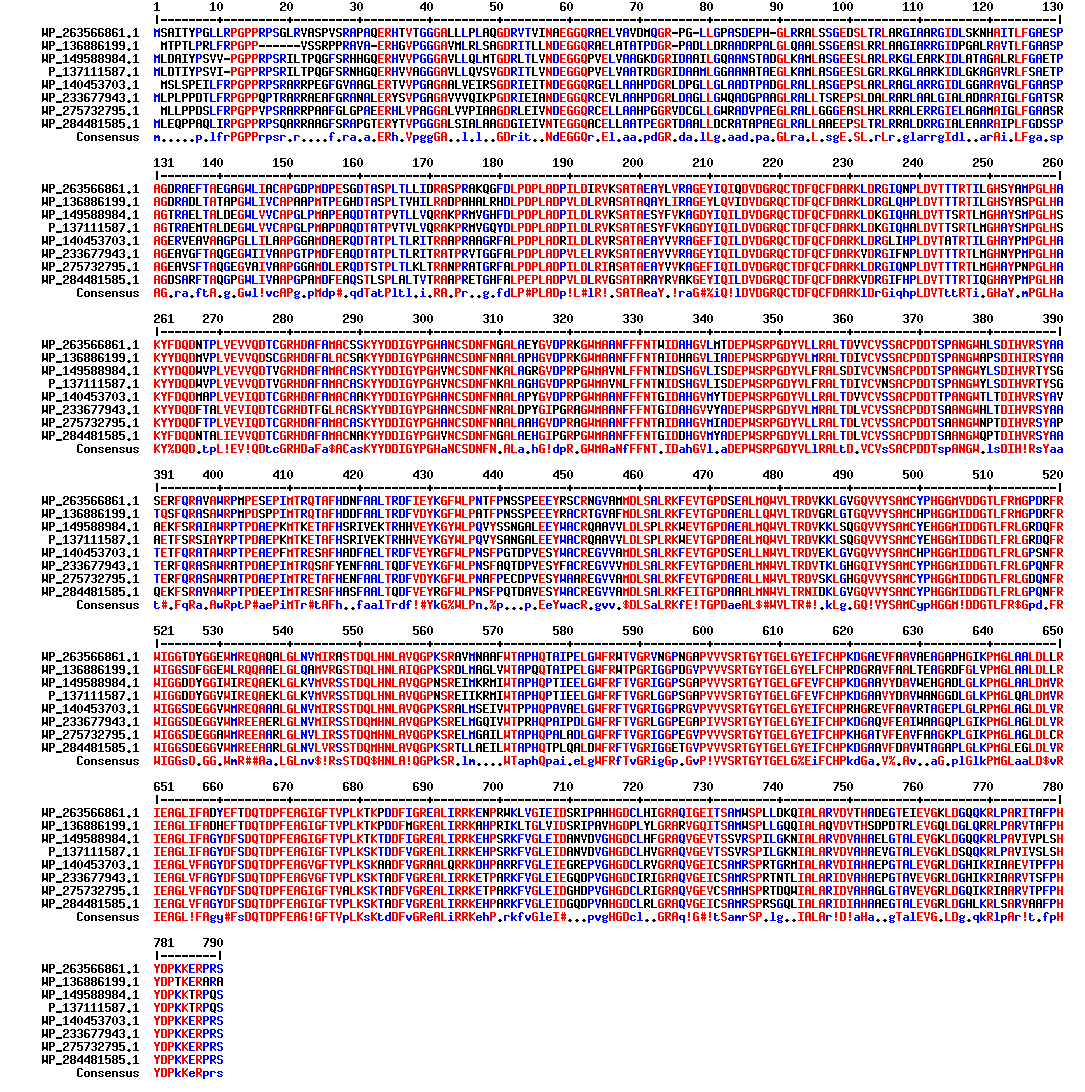

Figure S4. Sequence alignment of TDM from *Paracoccus* sp. and other organisms using *multalign*. Green arrows indicate key residues in active site binding to TMAO substrate. Red, conserved; blue, moderately conserved; black, no conserved residues. *Paracoccus* sp. DMF (WP_263566861.1), *Paracoccus gahaiensis* (WP_136886199.1), *Tabrizicola flagellates* (WP_149588984.1), *Rhodobacter* sp. SY28-1 (WP_137111587.1), *Amaricoccus solimangrovi* (WP_140453703.1), *Sinirhodobacter* sp. WL0062 (WP_233677943.1), *Paenirhodobacter* sp. CAU 1674 (WP_275732795.1), *Pseudodonghicola flavimaris* (WP_284481585.1).

**Figure S5.** **Zinc association with the TDM protein.** (**A) Energy-dispersive X-ray spectroscopy (EDS) analysis was performed to detect the presence and spatial distribution of zinc (Zn) in the purified TDM protein sample. The black-and-white panel represents the overall protein structure, while the green overlay indicates the specific localization of Zn signals, suggesting Zn binding or association with the protein. (B) Quantitative analysis of metal content was conducted using inductively coupled plasma mass spectrometry (ICP-MS). The molar ratios of iron and zinc relative to the TDM protein concentration were determined. Measurements were performed on independently prepared protein samples (n=2).**

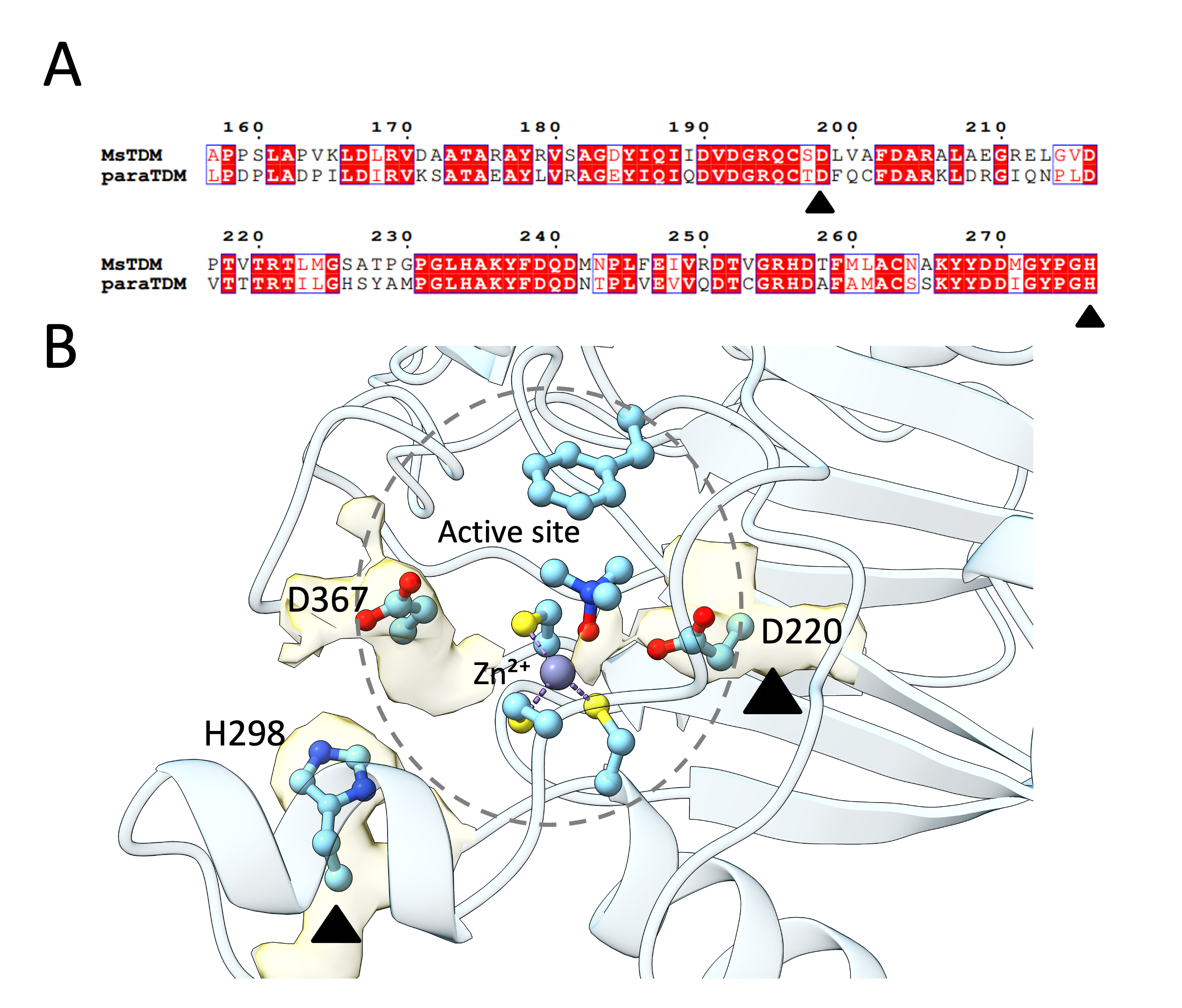

**Figure S6. paraTDM lacks an iron-binding site.** (A) Sequence alignment of *Methylocella silvestris* TDM (MsTDM, WP_012592556.1) and paraTDM (WP_263566861), highlighting residues proposed to coordinate Fe²⁺ in MsTDM (arrows). (B) Cryo-EM structure of TDM expressed in LB medium supplemented with 0.5 mM Fe(NH₄)₂SO₄ (Fe-enriched paraTDM). No additional EM density corresponding to a bound metal ion is observed, including in the regions between the proposed coordinating residues D220 and H298 (arrows) or between H298 and D367. Residues are shown in stick representation. The corresponding EM density contoured at 3.0 Å resolution are shown in the three residues.

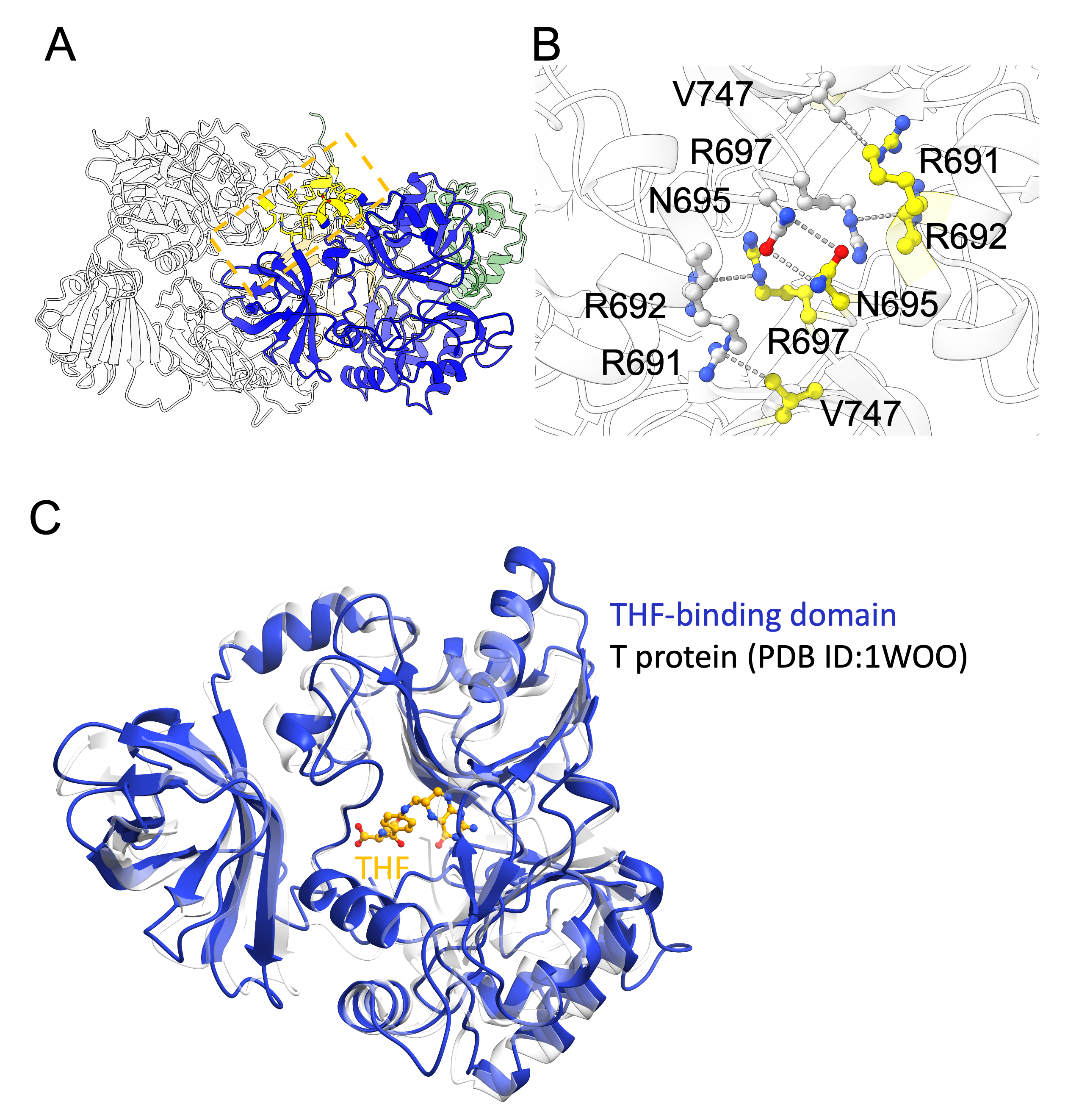

Figure S7. ****Structural features of the THF-binding domain in the TDM:DMA:HCHO:THF complex.**** (A) Dimerization of the THF-binding domains occurs through interactions between adjacent protomers, contributing to the overall oligomeric architecture of the TDM complex. (B) The central dimer interface is primarily formed by residues located on the inward-facing surfaces of the THF-binding domains. Stabilization of the interface involves multiple hydrogen bonds and additional polar contacts, facilitating a tight and specific interaction. (C) Structural comparison revealed a high degree of similarity to a THF-binding protein, as demonstrated by superposition with the T protein (PDB ID: 1WOO).

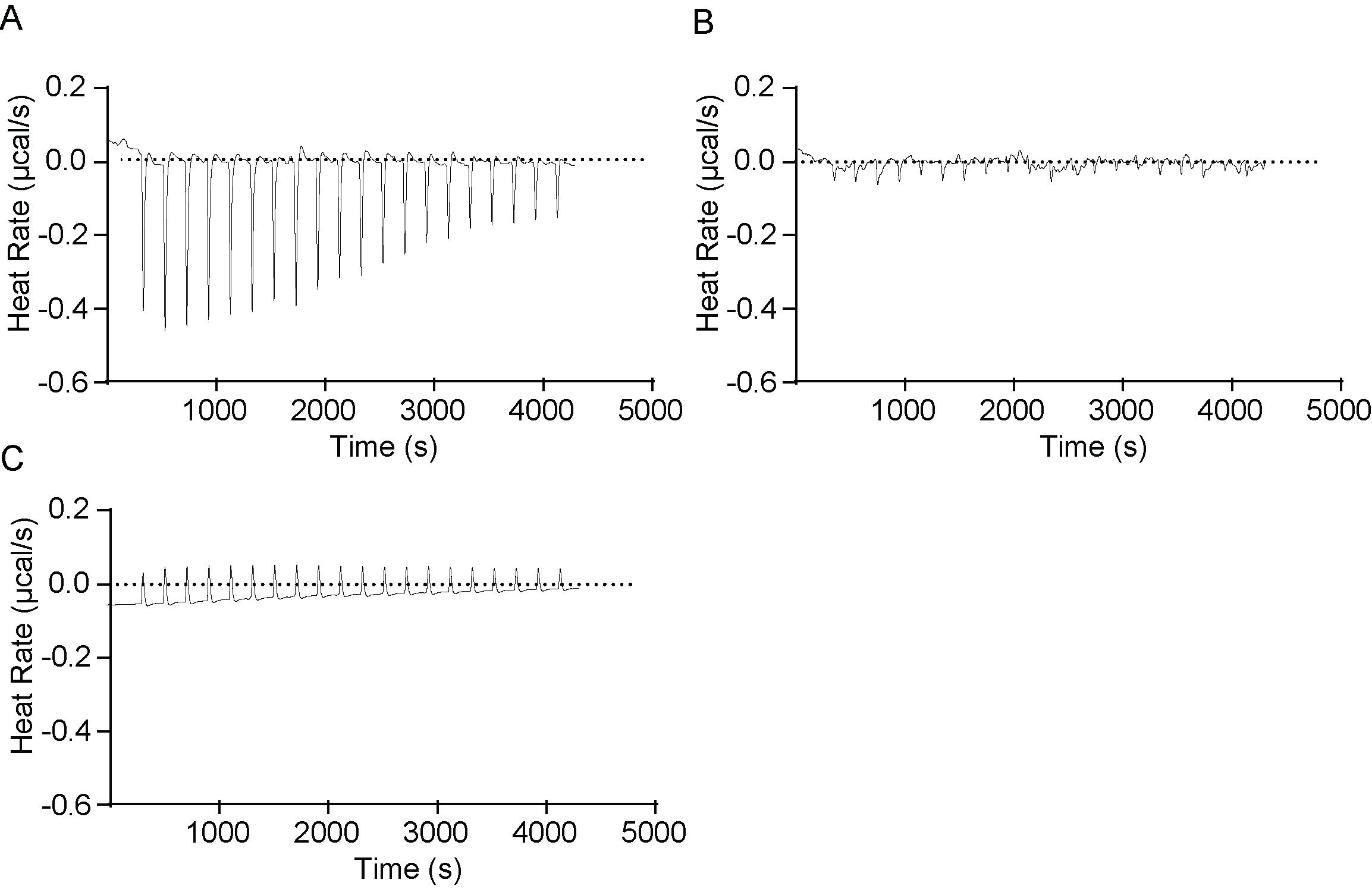

Figure S8. The binding of THF to TDM was determined by ITC. (A) Representative ITC thermogram showing the heat changes upon titration of THF into a solution containing purified TDM protein. (B, C) Control ITC thermogram showing titration of buffer into TDM (B), and titration of THF into buffer (C).

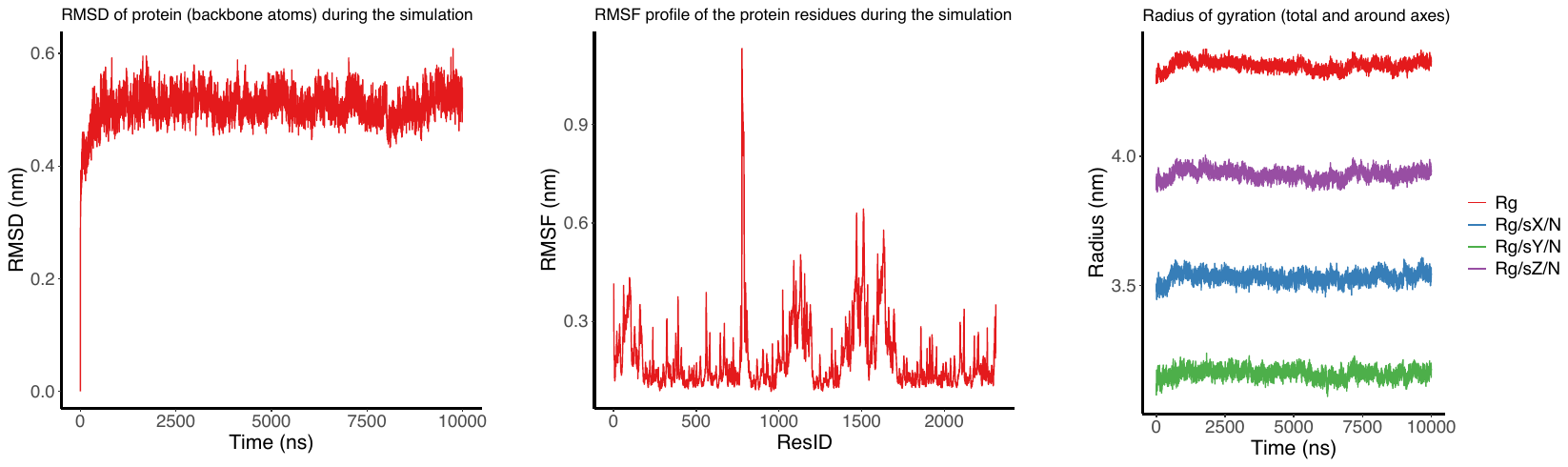

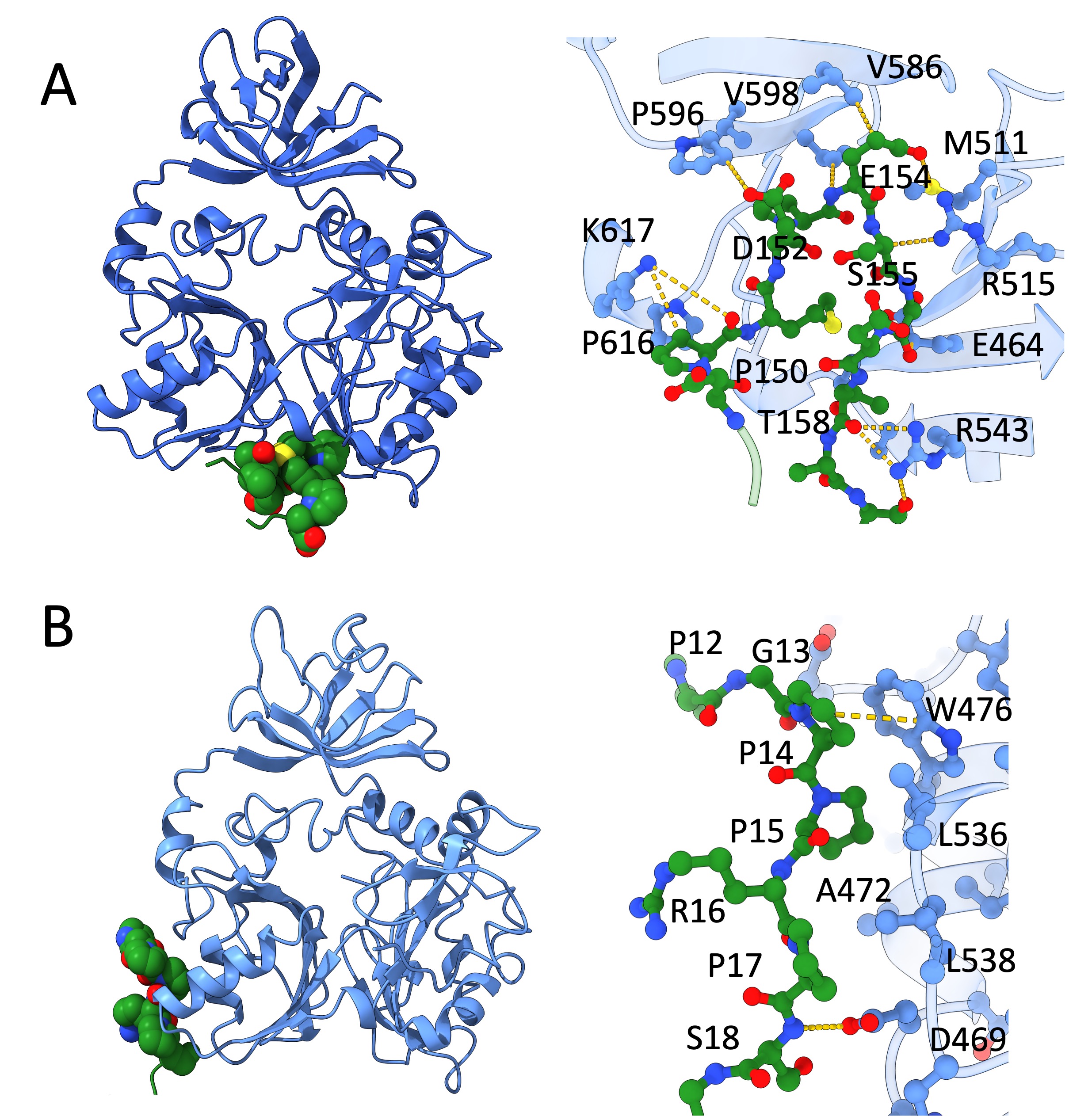

**Figure S9. Structural stability of the TDM complex during coarse-grained molecular dynamics (CG-MD) simulations.** Root-mean-square deviation (RMSD), root-mean-square fluctuation (RMSF), and radius of gyration (Rg) were analyzed over the course of a 10 μs CG-MD simulation. (A) The RMSD of protein backbone atoms relative to their initial positions stabilized around 0.5 nm, indicating overall structural stability. (B) The RMSF plot shows limited per-residue fluctuations, with elevated flexibility observed primarily in loop regions, consistent with expected dynamic behavior. (C) The Rg of the protein, calculated both overall and along individual spatial dimensions (x, y, and z), remained consistent throughout the simulation, further supporting the maintenance of a compact, folded structure.

****Figure S10.**** Two distinct binding modes between the N-terminal domain (green) and the THF-binding domains (blue and light blue). (A) Interaction interface between N-terminal residues 150–158 and the THF-binding domain. (B) Alternative interaction interface involving N-terminal residues 12–18 with the THF-binding domain.
